# Heterogeneity of the Frequency Domain Patterns in Persistent Atrial Fibrillation

**DOI:** 10.1101/285031

**Authors:** Shahriar Iravanian, Jonathan J Langberg

**Author notes:** Contact Information: Jonathan J. Langberg, M.D., Professor of Medicine, Emory University Hospital, 1364 Clifton Road, NE, Ste D403C, Atlanta, GA 30322.

## Abstract

**Background:** Persistent atrial fibrillation (AF) has remained a challenging clinical problem. The mechanisms of persistent AF are still subject to debate. Both a single mother-rotor with fibrillatory conduction and multiple meandering spiral waves have been proposed to explain persistent AF. Previous frequency domain studies have reported the presence of dominant frequency (DF) gradient (a marker of single mother-rotor) in paroxysmal, but not persistent AF.

**Methods and Results:** We performed temporally-dense high-resolution frequency domain analysis of 10-40 minutes segments of intracardiac signals recorded in 24 patients undergoing ablation of persistent AF. We observed two predominant patterns. The expected signature of the mother-rotor mechanism was observed in 38% of the patients. The frequency pattern in 54% consisted of two or more distinct frequency peaks with no obvious gradient, which is consistent with multiple separate primary spiral waves in electrophysiologically heterogeneous areas of atria. The average measured number of rotors per case was 1.71 ± 0.32, which provides a lower limit on the actual number of rotors. The single-zone pattern was exclusively seen in patients who were on a membrane-active antiarrhythmic medication at the time of ablation (P < 0.005).

**Conclusions:** AF is a heterogeneous disorder. High-frequency resolution analysis is a useful tool to detect the underlying mechanisms of AF and to classify it into patterns consistent with a single mother-rotor vs. multiple meandering wavelets.

## Introduction

Atrial fibrillation (AF) is the most common clinical arrhythmia and is responsible for significant morbidity and mortality. AF management has remained a challenging problem. Antiarrhythmic medications and ablation procedures are commonly used to control AF,^1^ but neither intervention is very effective, especially when AF progresses to the persistent stage.

Paroxysmal AF is commonly initiated by triggers from pulmonary veins.^2^ However, as AF progresses gradually to the persistent phase, the role of pulmonary veins diminishes and AF is sustained as a result of spiral waves and spatiotemporal chaos. The exact spatiotemporal organization of the spiral waves (rotors) and their response to various interventions (in the form of medications, ablation, or pacing) are not fully characterized and are still subject to intense debate.^3^ In particular, it is not clear whether AF is caused by a single anchored mother rotor with fibrillatory conduction to the rest of atria or whether it is the result of multiple interacting spiral waves with no obvious hierarchy (i.e., each spiral wave is as primary as the others). Settling this question has important ramifications in the management of AF. However, the complexity of AF has precluded an easy answer.

The pioneering works of Moe and later Allessie et al. let to the concept of multi-wavelet reentry as the mechanism of AF.^4–6^ Experimental work by Jalife and colleagues in late 1990s demonstrated the presence of a gradient in dominant frequency (DF) of intracardiac signals during AF, suggesting an alternative explanation of a single “mother” rotor, usually anchored in the posterior left atrium, with fibrillatory conduction to the rest of the atria.^7–9^ This hypothesis is supported (at least in some experimental models) by the direct visualization of stable spiral waves using optical mapping.^10^ Applying frequency-domain analysis to clinical AF in humans, Lazar et al. detected a DF gradient in paroxysmal but not persistent AF.^11^

Previously, we analyzed the intracardiac recordings during ablation of persistent AF using both time and frequency domain analysis methods to gain mechanistic insight into AF.^12,13^ We showed that temporally-dense and high-resolution frequency domain analysis is a valuable tool in elucidating the spatiotemporal dynamics of AF. Our main finding was that transitions from irregular and chaotic AF to regular flutter occur suddenly and have the hallmarks of phase transitions in complex dynamical systems.^14^

In this study, we expand on our previous work by focusing our attention on the dynamics and number of rotors during persistent AF. We postulate that it is possible to differentiate between the two putative mechanisms of AF by looking for their expected frequency domain signatures. In the case of a single rotor, we expect to see an area of fast and regular activity (a narrow peak in the frequency domain) with fibrillatory conduction to the rest of atria. This is the *classic* DF gradient pattern seen in previous studies. On the other hand, the predicted pattern for multiple independent rotors is different than DF gradient and is marked by multiple spatially-localized frequency peaks.

## Methods

### Patient Population and Ablation Procedure

The procedural and data collection methods were previously described.^12^ The study protocol was approved by the Emory University Institutional Review Board. The data were collected retrospectively from catheter ablation procedures for persistent AF done at the Emory University Hospital, Atlanta, GA. The procedure included radiofrequency-based isolation of pulmonary veins and substrate modification by targeting complex fractionated atrial electrograms (CFAE).^15^

All patients were in persistent AF at the time of ablation. A duodecapolar catheter (Livewire, St Jude Medical) was placed in the right atrium and advanced into the coronary sinus, such that on average, the distal five electrode pairs were in the coronary sinus and recording from the left atrium. Bipolar intracardiac signals from the duodecapolar catheter were recorded continuously for the duration of the procedure using a Cardio Lab System (GE Medical). The signals were band-passed filtered at 30-500 Hz, digitalized at 12 bits of resolution, sampled at 977 Hz, and were downloaded for offline processing.

### Signal Processing

Each channel was divided into 16384-point segments (∼16 seconds) with 50% overlap between them. The segments were detrended, rectified and transformed into the frequency domain with a resolution of 1/16 Hz. The resulting power spectra were normalized to the peak value at each segment. Short-Time Fourier Transform (STFT) graphs and the combined power spectrum of selected regions were used to describe the dynamics and were correlated with the raw intracardiac electrograms.

## Results

Episodes of AF in 24 patients undergoing ablation of persistent AF were classified into three groups based on the frequency domain analysis of a 10-40 min segment of uninterrupted AF:

1. **Single Zone:** in this group, there was a continuous frequency gradient from a fast area to the rest of the channels. The main signature of this type as seen in the power spectrum is a narrow and fast frequency peak with gradual slowing and widening in more distant channels (Figure 1A).

**Figure 1.**
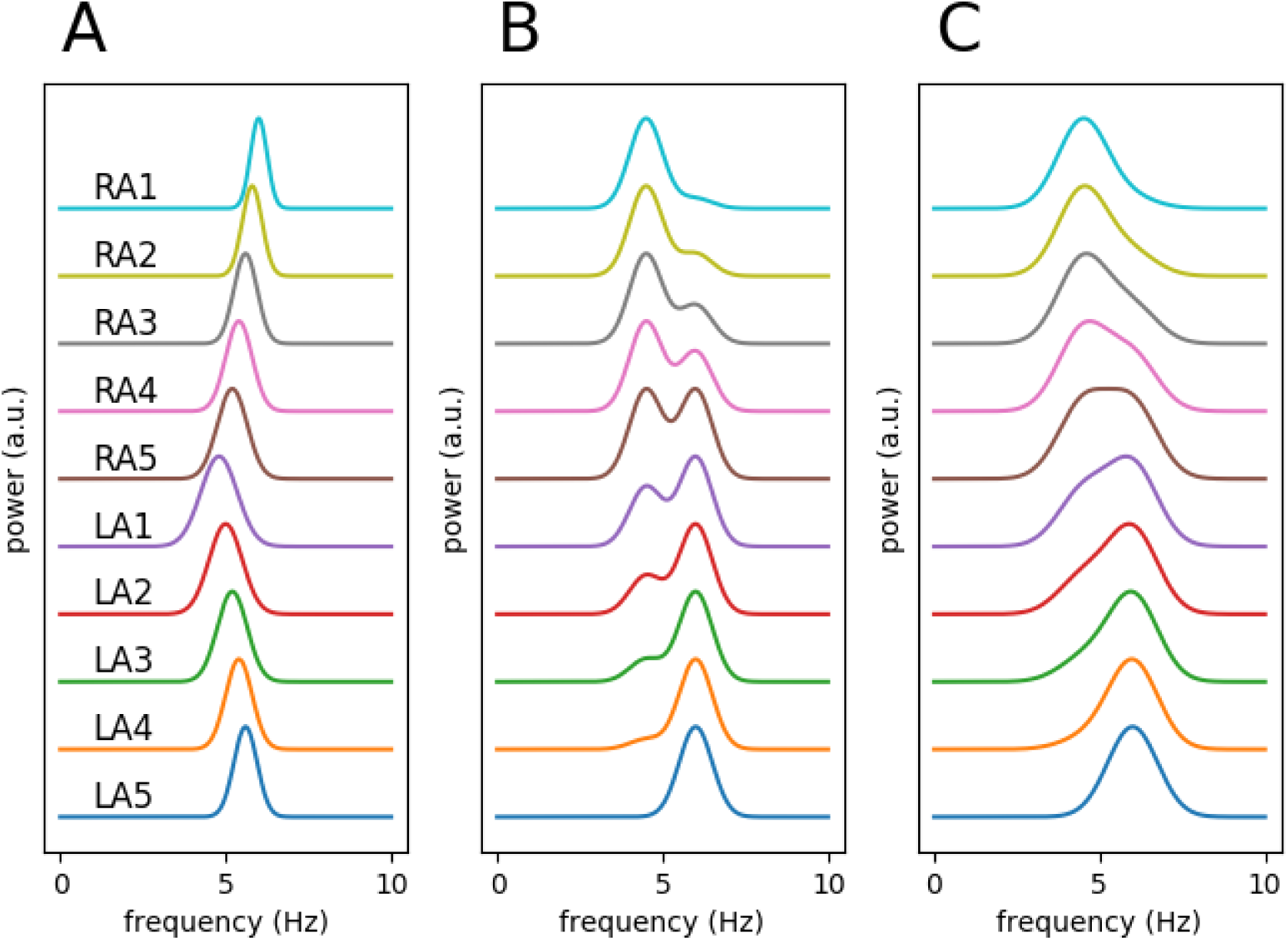
Schematics of different frequency patterns. **A** shows a single-zone pattern with a narrow peak at the fastest frequency (RA1) and a frequency gradient and wider peaks toward the slower frequencies. The resulting pattern is ‘‘chevron-shaped”. **B** displays a dual-zone pattern. **C** is the same data as **B**, but depicted at a lower frequency resolution. Fusion of the peaks, especially in the middle spectra, results in an apparent gradient and may be interpreted by mistake as a single-zone.
2. **Dual- or Multi-Zone:** in this group, there were two or more distinct frequency zones with no apparent gradient. In some cases, there were overlaps between the zones, such that a single channel straddled more than one zone and exhibited two or more peaks in the power spectrum (Figure 1B). It should be noted that if we had used a shorter Fourier Transformation window (e.g., a 2 to 4-second window instead of the 16-second one), the resulting lower resolution would have smeared and obscured the dual peaks and the resulting pattern could have been misidentified as a single zone (Figure 1C).
3. **Complex Pattern:** this is a mixture of the two previous types with multiple distinct zones with areas of frequency gradient.

The mean age of the patients was 61 years old (95% confidence interval [CI] 45-77 years). Fifteen patients were male (63%).

We observed the single zone type in 9 out of the 24 cases (38%). The multi-zone signature was seen in 13 cases (54%, 11 with two zones and 2 with three or more). The frequency patterns were too complex to distinguish between the single or multi-zones in two cases and were therefore designated as complex. The average number of detected rotors per case was 1.71 (CI 1.39-2.03). This is a lower bound on the actual number of rotors as the duodecapolar catheter samples only a portion of the atria.

Fourteen patients were taking a class I and class III antiarrhythmic medication at the time of ablation, 9 exhibited a dual/multi-zone pattern, and 5 displayed a single-zone pattern. On the other hand, all the patients who were not on an antiarrhythmic medication had a dual/multi-zone pattern (P < 0.005, Fisher’s exact test).

Figure 2 displays a clear cut example of a single zone with frequency gradient. In this case, the right atrium is in a stable flutter, but the left atrium is in AF. There is a single narrow peak on the right atrial channels in the power spectra (panel C) with a gradual slowing and widening as we move toward the proximal coronary sinus. We interpret this pattern as a flutter in the right atrium with fibrillatory conduction to the left atrium through the Bachmann’s bundle.

**Figure 2.**
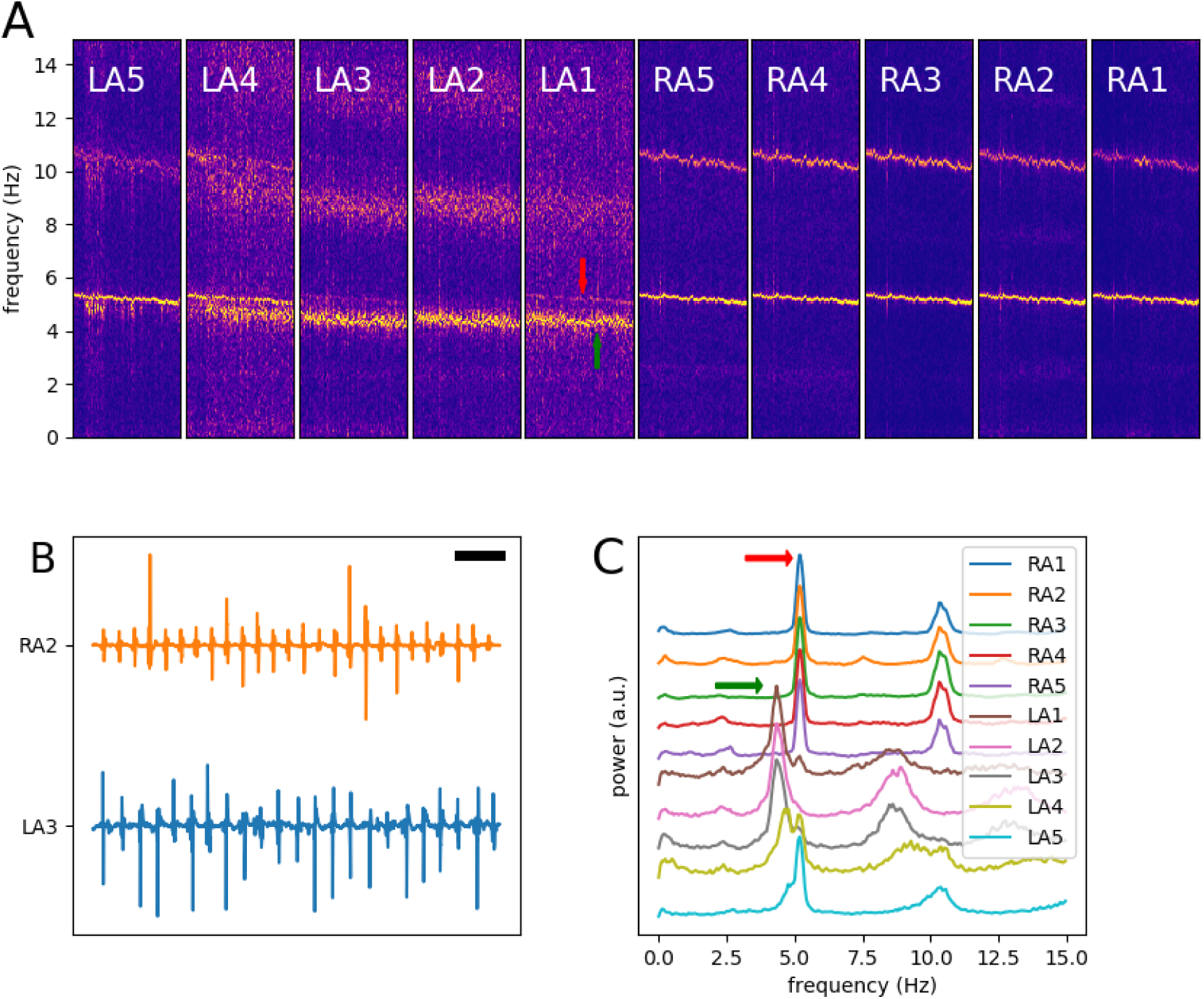
An extreme example of single-zone frequency pattern. **A** depicts the STFT spectra of a 15-minute segment of intracardiac signals obtained from ten recording channels during AF. The right atrial (RA1-RA5) channels have a narrow spectral peak, consistent with flutter. The left atrial (LA1-LA5) channels show double peaks: the same narrow signal as the left (red arrow), and a wider and slower peak correlated with fibrillatory conduction (green arrow). **B** Two 5-second segments of unprocessed intracardiac signals. The black horizontal bar is 500 ms. **C** The averaged spectra. The spectra are shifted vertically for better visualization.

Most cases of single zone type are not as dramatic as Figure 2. Figure 3 depicts a typical single zone case. Here, the peak frequency is in the lateral right atrium with a gradual slowing toward the proximal coronary sinus, as seen by the chevron pattern of the peaks in the spectrograms (similar to Figure 1A).

**Figure 3.**
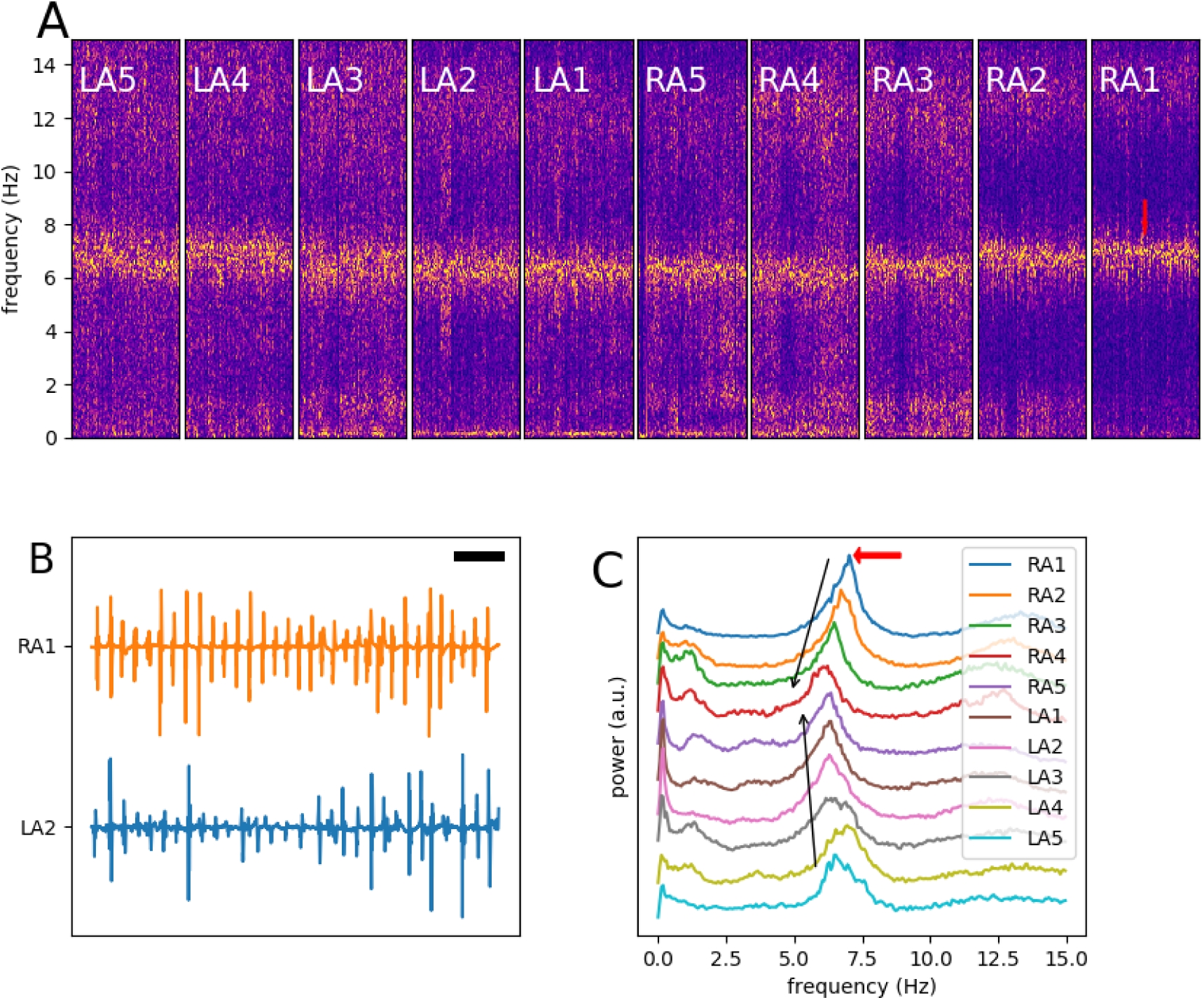
A typical example of single-zone frequency pattern. **A** depicts the STFT spectra of a 30-minute segment of intracardiac signals obtained from ten recording channels during AF. The fastest channel with a narrow frequency peak is RA1 (red arrow). There is gradual widening and slowing of the peak toward LA1/RA5 region. **B** Two 5-second segments of unprocessed intracardiac signals. Note regular activity in RA1 and slower and fractioned atrial signals in LA2. The black horizontal bar is 500 ms. **C** The averaged spectra. The spectra are shifted vertically for better visualization.

Figure 4 represents a typical dual zone AF. There are two distinct and widely separated frequency peaks: a fast one at 6.0 Hz, which is mainly seen in the left atrial channels, and a slower predominantly right-atrial peak at 4.5 Hz. These peaks remain well separated with no gradual transition from one to the other. The intracardiac signals (panel B) from the two zones show regular activity with different cycle lengths. This pattern is in contrast to Figure 3B, where there is a fractioned signal on the slower side. Another dual zone example is presented in Figure 5. Again, there are two distinct peaks. However, their separation is less (5.3 and 6.3 Hz), and the two peaks are partially fused in the middle channels (similar to Figure 1C).

**Figure 4.**
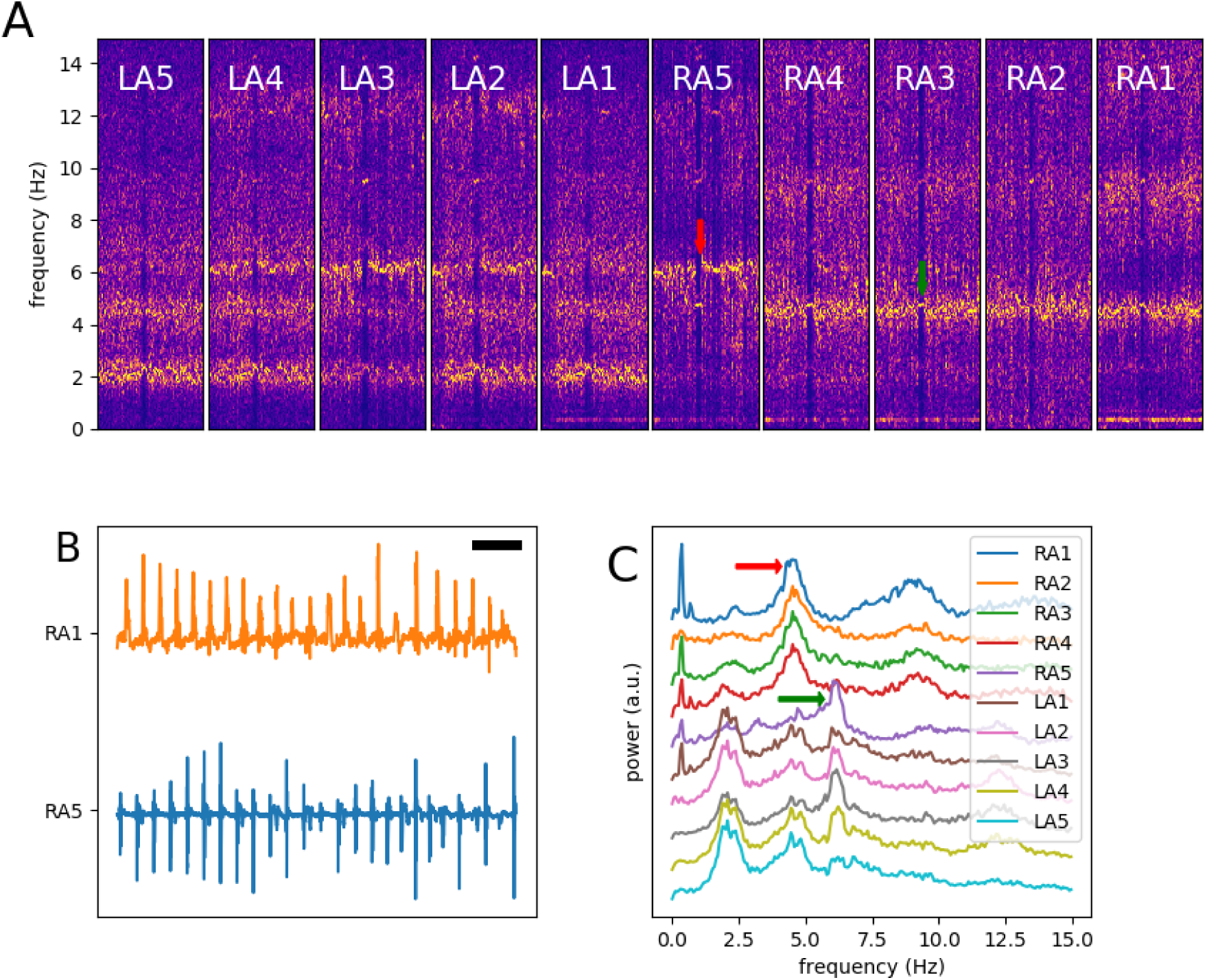
A typical example of dual-zone frequency pattern. **A** depicts the STFT spectra of a 10-minute segment of intracardiac signals obtained from ten recording channels during AF. There are two well-separated frequency peaks (red and green arrows). Channels LA1 to LA4 exhibit both peaks. **B** Two 5-second segments of unprocessed intracardiac signals. The black horizontal bar is 500 ms. **C** The averaged spectra. The spectra are shifted vertically for better visualization.

**Figure 5.**
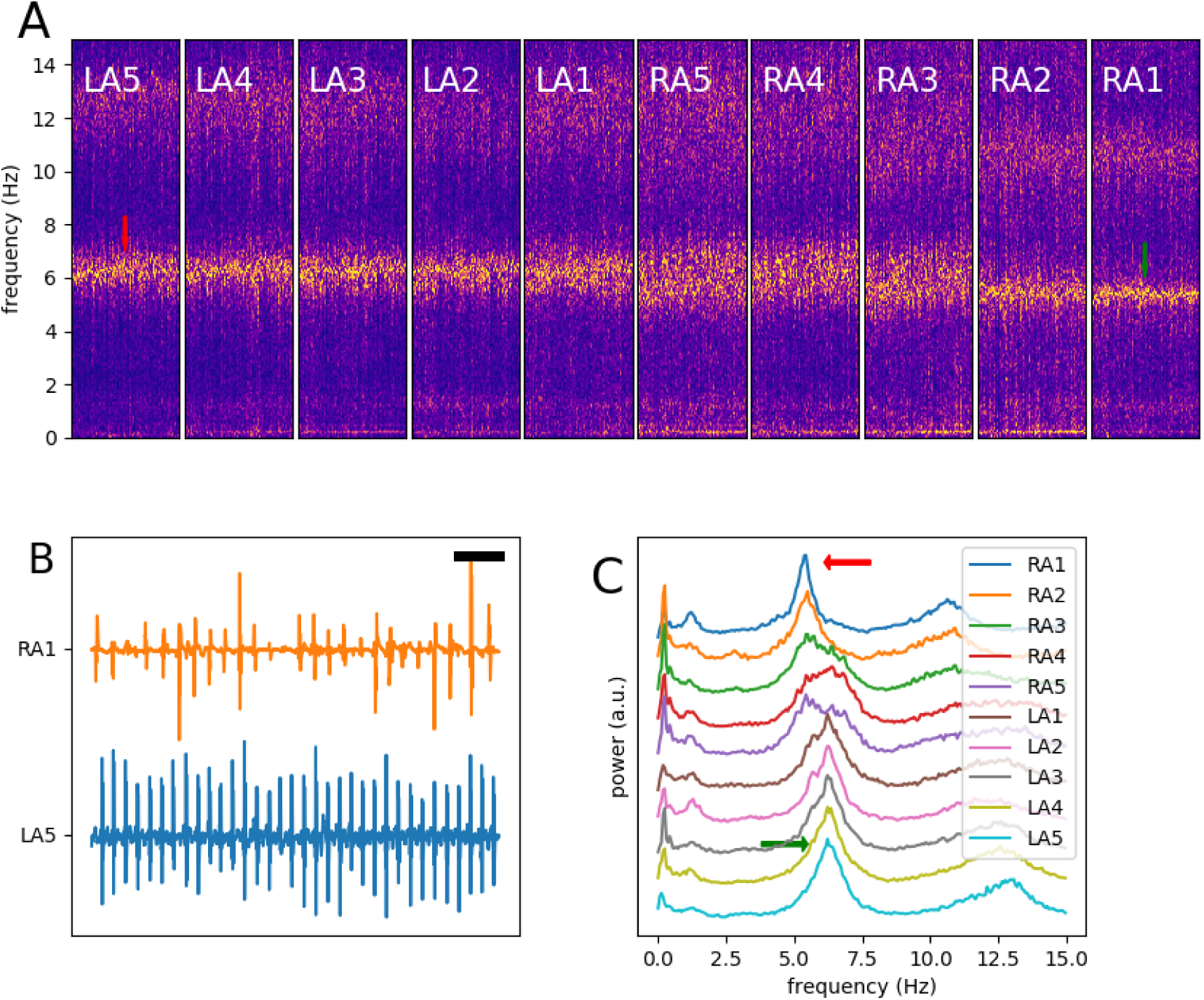
Another example of dual-zone frequency pattern. **A** depicts the STFT spectra of a 20-minute segment of intracardiac signals obtained from ten recording channels during AF. There are two frequency peaks (red and green arrows). The separation between the channels is less than Figure 4. Channels RA3 to RA5 exhibit fused frequency peaks. **B** Two 5-second segments of unprocessed intracardiac signals. Note fast and regular activity in LA5. The black horizontal bar is 500 ms. **C** The averaged spectra. The spectra are shifted vertically for better visualization.

Some cases exhibit more than two peaks. Figure 6 is an example of at least three distinct frequency peaks. The slowest peak (seen mainly in channel RA5) is very narrow, suggestive of regular activity. Interestingly, there is also a hint of a fourth peak in channel LA5, which is fast and narrow.

**Figure 6.**
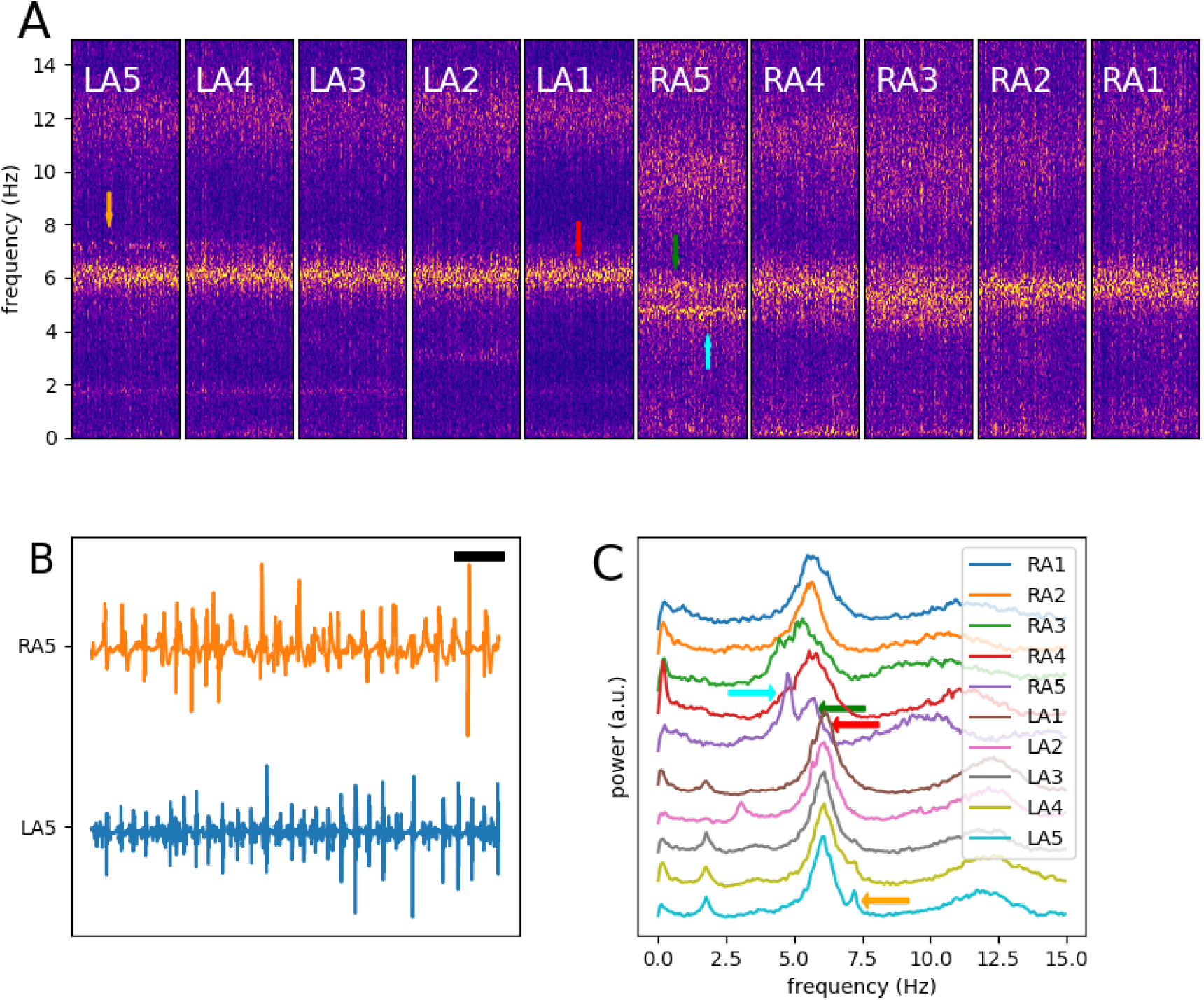
An example of multi-zone frequency pattern. **A** depicts the STFT spectra of a 20-minute segment of intracardiac signals obtained from ten recording channels during AF. There are four frequency peaks (red, green, yellow and cyan arrows). Multiple channels show double peaks: red/yellow in LA4 and LA5, green/cyan in RA3 and RA5, and possibly red/green in RA1. **B** Two 5-second segments of unprocessed intracardiac signals. Note fast and irregular activity as a result of superimposition of multiple peaks. The black horizontal bar is 500 ms. **C** The averaged spectra. The spectra are shifted vertically for better visualization. Note the presence of two well-defined frequency peaks that can be followed from the beginning of the recording up to the transition at 167 min. The potassium-channel blocker ibutilide was infused at 160 min, resulting in the global slowing of atrial activities. **B** and **C** show partial STFT of channels LA4 and RA4 from Figure 5. Note that the frequency peak in **B** is lower. Also, the flutter emerges from the slower peak (**B**) rather than the faster peak in **C**.

### Pattern Stability

In this study, the frequency patterns were assigned based on the data recorded while ablation lesions were applied. To show the stability of patterns during AF, we divided the period of AF from the beginning of the ablation to the time of the first transition to a flutter or sinus rhythm into two halves and compared the peaks in the frequency domain. Figure 7 shows four representative examples. In some cases, the frequency peaks shifted slightly toward lower frequencies as a result of ablation, but the relative structure of the peaks and the designation to a single or multi-zone pattern remained the same.

**Figure 7.**
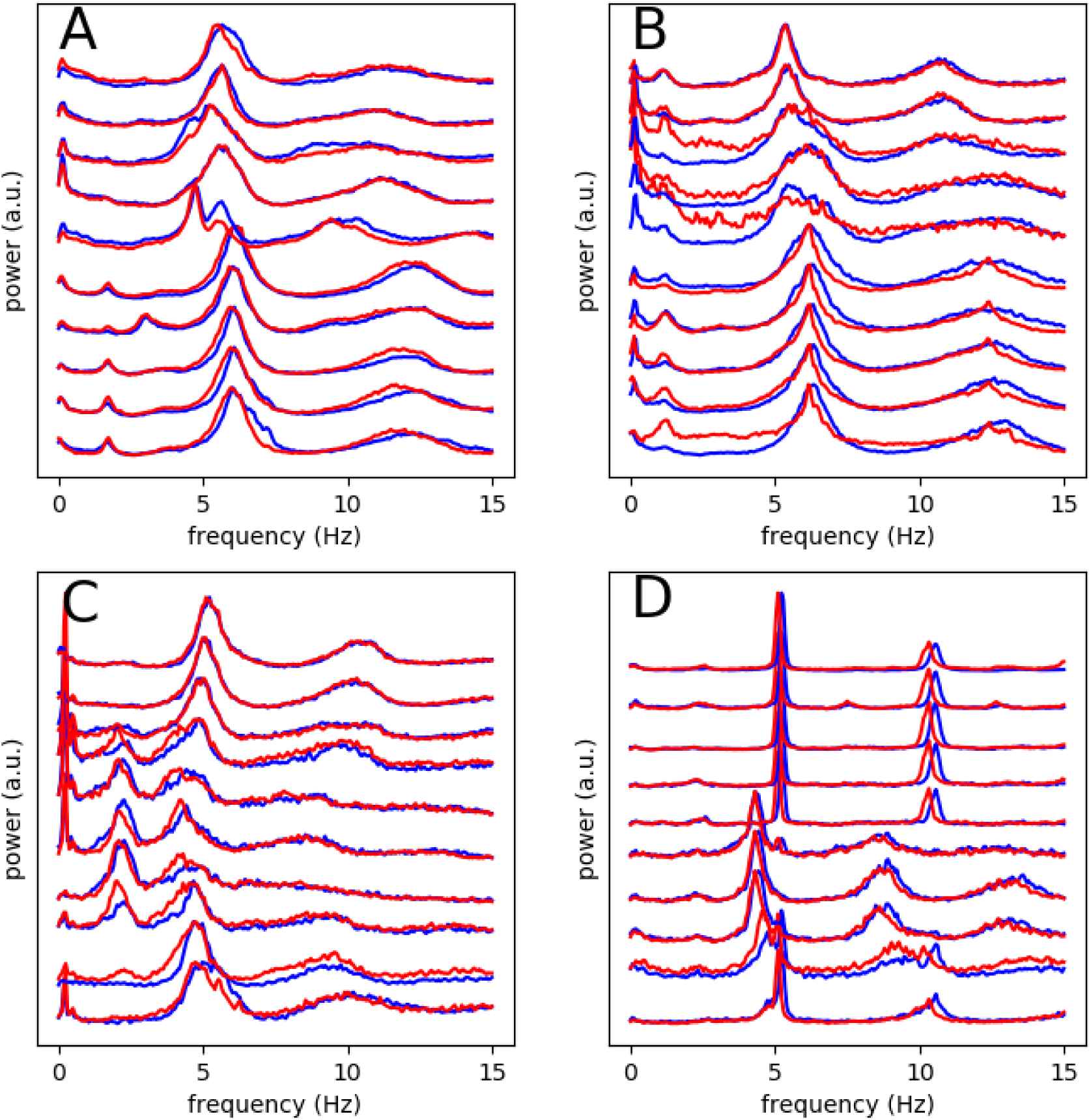
Comparison of frequency patterns during AF ablation. AF episodes from the beginning of ablation to the time of the first termination were divided into two halves. Each panel displays the overlapped frequency peaks. The first half is drawn in blue and the second half in red. Note the near perfect match between the two halves. The main difference is that the second halves are slightly shifted toward lower frequencies, but the general pattern remains the same. **A** and **B** show a multi-zone pattern, and **C** and **D** a single-zone pattern.

### Transitions from AF to Flutter

Previously, we have described sudden transitions from AF to a regular flutter.^12,13^ A detailed analysis of these transitions is useful in revealing the underlying dynamics. Figure 8A shows such a transition in a multi-zone AF. In this case, ablation per se did not organize the rhythm to a flutter, and ibutilide was infused near the end of the case to promote the transition. Before the ablation, two distinct peaks were seen throughout the case. After ibutilide infusion, both peaks slowed down and then flutter emerges from the lower frequency peak, as if the eventual flutter was present, but constrained to one region, during persistent AF. This finding argues that the slower peak is also a primary reentrant source and not merely a manifestation of conduction block from a faster source.

**Figure 8.**
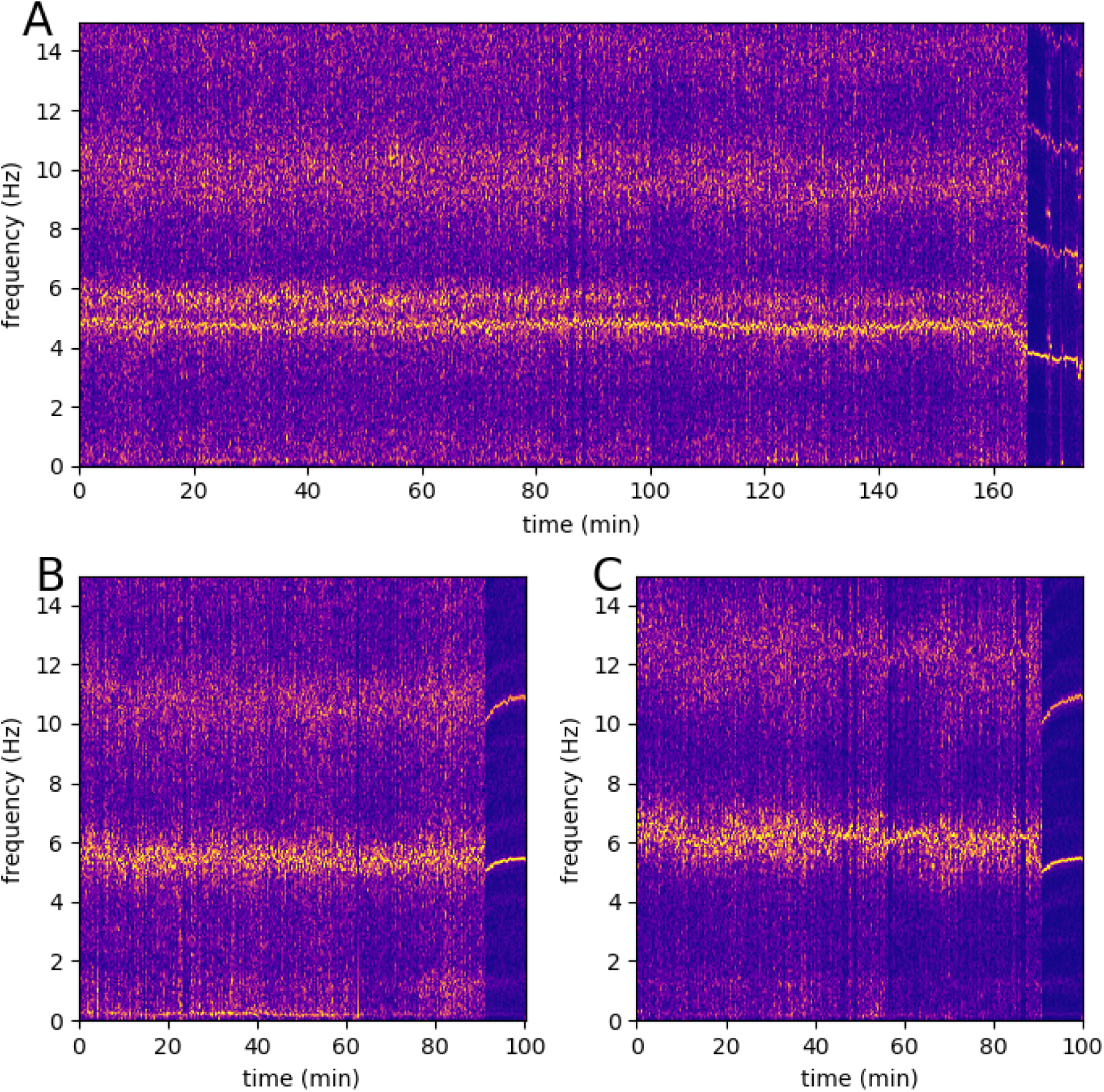
Examples of AF to flutter transitions. **A** shows the full STFT of channel RA5 from Figure 6.

A similar pattern is present in Figure 8. Panels B and C, corresponding to different channels of a dual zone case, follow the slower and faster peaks, respectively. Ablation-induced flutter emerges from the slower peak.

## Discussion

AF is a heterogeneous disease. The mechanisms of AF may vary in different patients and can evolve and changes over time in the same individual as a result of electrical and structural remodeling.^16^ This heterogeneity of mechanism manifests as variable frequency domain patterns of intracardiac signals. However, in this study, we show that the frequency spectra during persistent AF are stable and can be broadly classified into two major groups: single-zone and multi-zone patterns.

We detected the frequency domain signature of a single rotor in less than 40% of the cases. The majority of the rest are classified as dual or multi-zone patterns, which we interpret as the evidence of multiple distinct *driver domains* in electrophysiologically heterogeneous regions of atria,^17^ and not necessarily multiple coexisting anchored and localized spiral waves. Therefore, both the mother rotor and multiple wavelet hypotheses may be applicable in certain subgroups of patients with persistent AF.

The recognition of single vs. multi-zone pattern in a given patient may have therapeutic implications. For example, Focal Impulse and Rotor Modulation (FIRM) technique was initially developed assuming a single mother rotor as the underlying mechanism of AF.^18^ However, clinical studies did not corroborate this assumption. Later clinical studies using FIRM reported an average 2.8 ± 1.4 source per patient.^19^ Based on our results, it is possible that the group of patients who have in fact a single rotor are better suited for FIRM ablation.

Membrane-active antiarrhythmic medications (class I and III) had a significant effect on the frequency pattern of AF. Specially, we observed no single-zone pattern in patients who were not taking an antiarrhythmic. These medications, by reducing the excitability (for sodium-channel blockers) or prolonging the action potential duration (for the potassium-channel blockers), increase the reentry cycle length and decrease the potential number of rotors that can be fit onto atria. This result may explain the failure to detect DF gradient in persistent AF in previous studies,^11^ and argues for performing rotor mapping without interruption of antiarrhythmic medications at the time of the procedure.

Temporally-dense frequency domain analysis is a useful tool that can be used to classify AF before interventions such as ablation. As is evident from visual inspection of the unprocessed signals in Figures 2 to 6, it is not possible to discern the patterns which are readily apparent in the frequency domain. Moreover, this kind of analysis can be helpful in characterizing and localizing rotors, especially if applied to multi-channel basket catheters covering a larger area of atria. For example, in the single-zone with gradient pattern, the anchored spiral wave is likely to be closer to the channel with a fast and narrow frequency peak, whereas this statement is not necessarily true for the multi-zone pattern. Also, it is plausible that the underlying pattern is correlated with the immediate and long-term results of ablation.

From a practical standpoint, the ability to resolve frequency peaks requires a minimum frequency resolution of ∼0.1 Hz (0.067 Hz in our study). Many dominant frequency (DF) studies in AF employ a window too short for robust detection of the different patterns.

## Limitations

Our data was collected during ablation of persistent AF. The ongoing ablation affects the frequency content of the signals. However, the pattern remained stable during ablation.

The relatively small sample size of the study precludes the assessment of the association between the frequency patterns and the success of ablation.

The interpretation of the different patterns as signifying a single or multiple rotors is indirect, as the recording was done by a duodecapolar catheter placed in the right atrium and coronary sinus. This provides only a limited and far from the action view of the atria (spiral waves preferentially localize to the posterior left atrium and areas around the pulmonary veins). Also, we expect that such a limited view underestimates the actual number of rotors.

